# Functional constraints on replacing an essential gene with its ancient and modern homologs

**DOI:** 10.1101/087924

**Authors:** Betül Kacar, Eva Garmendia, Nurcan Tunçbağ, Dan I. Andersson, Diarmaid Hughes

**Author notes:** Equal contributors (Betül Kacar, Eva Garmendia). Address correspondence to: Betül Kacar.

## Abstract

The complexity hypothesis posits that network connectivity and protein function are two important determinants of how a gene adapts to and functions in a foreign genome. Genes encoding proteins that carry out essential informational tasks in the cell, in particular where multiple interaction partners are involved, are less likely to be transferable to a foreign organism. Here we investigated the constraints on transfer of a gene encoding a highly conserved informational protein, translation elongation factor Tu (EF-Tu), by systematically replacing the endogenous *tufA* gene in the *Escherichia coli* genome with its extant and ancestral homologs. The extant homologs represented *tuf* variants from both near and distant homologous organisms. The ancestral homologs represented phylogenetically resurrected *tuf* sequences dating from 0.7 to 3.6 bya. Our results demonstrate that all of the foreign *tuf* genes are transferable to the *E. coli* genome, provided that an additional copy of the EF-Tu gene, *tufB*, remains present in the *E. coli* genome. However, when the *tufB* gene was removed, only the variants obtained from the γ-proteobacterial family (extant and ancestral), supported growth. This demonstrates the limited functional interchangability of *E. coli tuf* with its homologs. Our data show a linear correlation between relative bacterial fitness and the evolutionary distance of the extant *tuf* homologs inserted into the *E. coli* genome. Our data and analysis also suggest that the functional conservation of protein activity, and its network interactivity, act to constrain the successful transfer of this essential gene into foreign bacteria.

## Introduction

The Complexity Hypothesis assigns the function and the network interactivity of a gene product as two primary factors determining a gene’s capacity to successfully transfer and adapt in another genome (Jain, et al. 1999; Sorek, et al. 2007; Wellner, et al. 2007; Cohen, et al. 2011; Kitahara and Miyazaki 2013). Consequently, genes involved in central informational tasks (i.e. replication, transcription and translation) are expected to be less transferable to a foreign genome, in comparison with genes involved in other activities (Smith, et al. 1992; Jain, et al. 2003; Sorek, et al. 2007).

Genomic replacement of an essential gene with another homolog may potentially disturb the gene product’s function depending on the degree of functional equivalence or compatibility between the two genes (Shen and Huang 1986; Kacar and Gaucher 2013; Blank, et al. 2014; Bershtein, et al. 2015). Alterations in gene dosage resulting from the acquisition of a foreign gene may also perturb cellular homeostasis resulting in a lower transcription rate or an altered global pattern of gene expression, network function and organismal survivability (Ito, et al. 1998; Coulomb, et al. 2005; Drummond, et al. 2005; Durfee, et al. 2008; Zotenko, et al. 2008; Lind and Andersson 2013; Bershtein, et al. 2015; Karcagi, et al. 2016).

Genes involved in the translation of mRNA into protein perform one of the most crucial informational tasks in the cell, and based on phylogenomic analysis they are expected to be highly resistant to gene transfer (Ciccarelli, et al. 2006; Sorek, et al. 2007). There are examples demonstrating that some ribosomal protein genes can be integrated into foreign genomes under certain conditions (Brochier, et al. 2000; Lind, Berg, et al. 2010; Lind, Tobin, et al. 2010; Knoppel, et al. 2016). However, few studies have systematically tested whether there is a direct correlation between organism fitness and the evolutionary distance between an essential endogenous gene and its substituted (ancestral or extant) homolog.

Here we focus on the bacterial elongation factor Tu (EF-Tu) protein, encoded by tuf, one of the most ancient and highly conserved proteins. EF-Tu has an essential function in the translation machinery by delivering aminoacylated tRNA molecules into the A-site of the ribosome (Miller and Weissbach 1977). EF-Tu is encoded by two genes in *E. coli, tufA* and *tufB*, generated by an ancient duplication event thought to be specific to the proteobacterial lineage preceding the Cambrian period (Baldauf, et al. 1996; Lathe and Bork 2001). Expression of EF-Tu is primarily driven by *tufA*, with 66% of cellular EF-Tu expressed from the *tufA* gene (Abdulkarim and Hughes 1996). EF-Tu protein levels in the cell are correlated with cellular fitness and intrinsically regulated in order to maintain growth rate (Tubulekas and Hughes 1993a; Brandis, et al. 2016; Kacar, et al. 2016). EF-Tu belongs to the ancient protein repertoire of the cell, evolves slowly and serves as a functional fossil by participating in ancient and conserved functions (Hsiao, et al. 2009). It remains unclear whether *tuf* genes are replaceable by their ancient counterparts or homologs obtained from an extant organism. Answering this question would allow us to explore the limits of interchangeability for the *E. coli tuf* gene, and to ascertain a pattern within and between bacterial lineages across time and divergence. We sought guidance from a methodology referred to as ancestral sequence reconstruction (Pauling and Zuckerkandl 1963; Benner 1995; Thornton 2004; Benner, et al. 2007; Gaucher, et al. 2008; Kacar 2016) and accessed reconstructed ancestral *tuf* variants, as well as modern *tuf* gene sequences in order to observe the patterns of interchangeability among multiple nodes along the EF-Tu phylogenetic tree. We utilized a set of foreign genes representing *E. coli* EF-Tu homologs from closely- and distantly-related bacteria, as well as phylogenetically inferred ancestral EF-Tu proteins dating from 0.7 to 3.6 bya, thus accessing interspecies (modern) and ancestral (paleogenetic) axes. We determined the fitness effects of the introduction of a foreign *tuf* gene into each strain by replacing the native *E. coli tufA* gene with foreign variants, and asked whether these foreign genes could support cell viability when the *tufB* gene was removed from the chromosome. We examined the impact of the *tuf* gene replacements on growth rate and protein levels, as well as on protein function-structure, and assessed the extent of lateral and ancestral phylogenetic distances between the alien gene and host genome that yielded viable organisms.

## Results

### Replacement of the *tufA* gene in *E. coli* reduces relative fitness

Using genetic recombineering, we generated a set of *E. coli* strains in which the *tufA* gene coding sequence was precisely replaced by the coding sequence of its ancestral and modern homologs (Figure 1, Figure S1A). The 16 *tuf* homologs cover bacterial species from a wide span of taxa *(Yersinia enterocolitica, Vibrio cholerae, Pseudomonas aeruginosa, Legionella pneumophila, Bartonella hanselae, Streptococcus pyogenes, Bacillus subtilis, Thermus thermophilus, Mycobacterium smegmatis* and *Thermatoga maritima)* as well as six ancestral sequences that extend deep into the bacterial phylogenetic tree (Figure 1). These six sequences represent ancestral nodes dating from approximately 0.7 bya, back to the last common ancestor of bacterial *tuf*, dated to approximately 3.6 bya (Battistuzzi, et al. 2004; Gaucher, et al. 2008). These homologs of *tuf* encode EF-Tu variants that range in amino acid identity, from 93.9% (Y. *enterocolitica)* to 69.4% *(T. maritima)*, relative to *E. coli* EF-Tu (Figure S2). Nucleotide sequences of the ancestral *tuf* genes are shown in Table S4.

**Figure 1.**
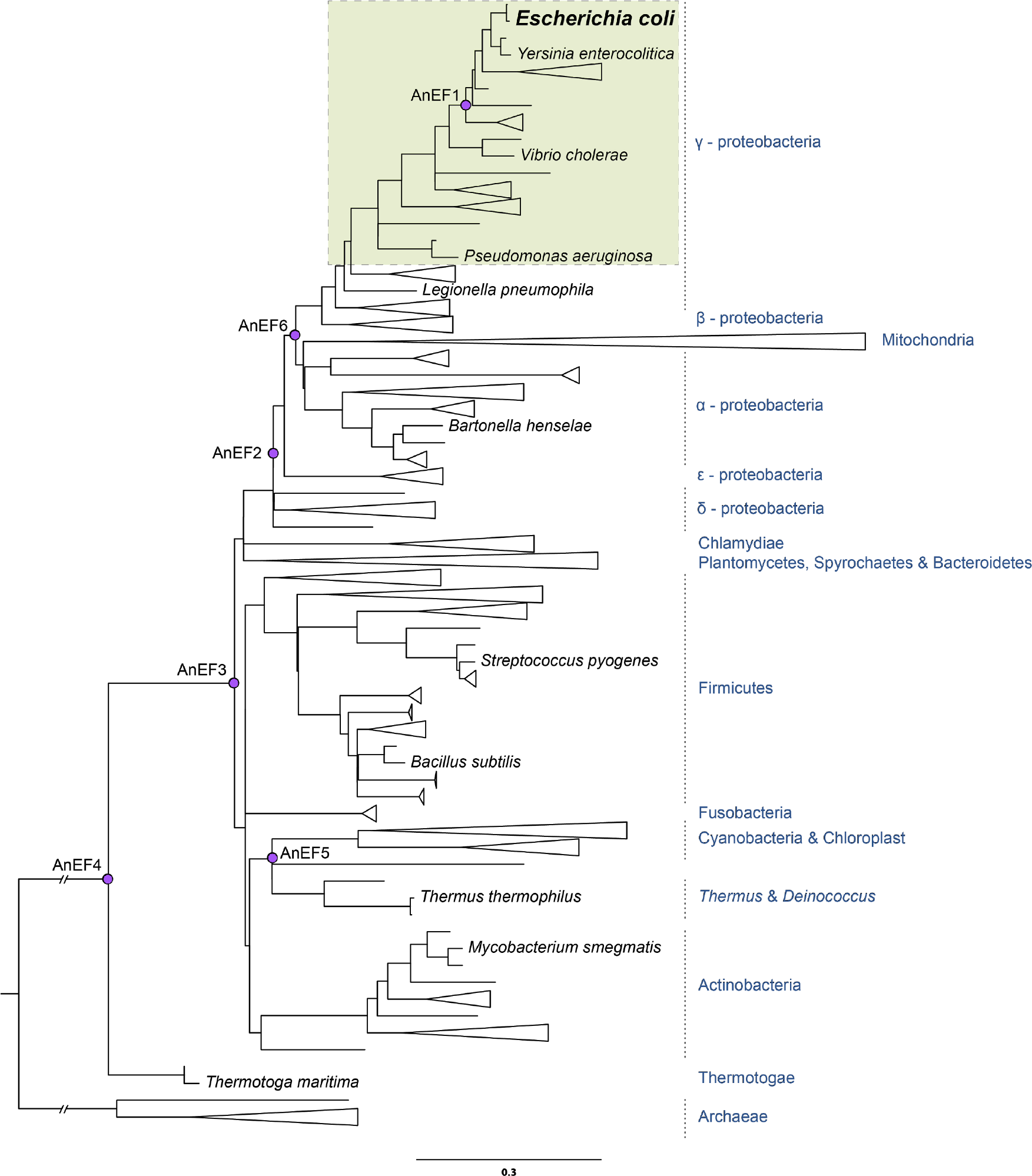
Phylogenetic tree indicating the node and taxa of the ancestral and modern *tuf* (EF-Tu) homologs. Pink circles represent the ancestral EF-Tu nodes. *E. coli* was genetically engineered to carry ancestral or modern homologs of tuf, encoding translation elongation factor EF-Tu, replacing the native *E. coli tufA* gene. A shaded box indicates the area of viability of EF Tu gene exchange. The scale bar expresses units of amino acid substitutions per site. The tree is adapted from (Gaucher et al. 2008, Battistuzzi et al. 2004).

To construct a set of isogenic strains, the endogenous *E. coli tufA* gene was replaced with each of the *tuf* variants while the second endogenous *tuf* gene, *tufB*, remained intact in the genome. All engineered strains in which *tufA* was replaced with a foreign *tuf* gene retained viability (Figure 2). The effects on relative fitness of the foreign *tuf* genes were determined by measuring the exponential growth rates in LB and relating them to that of the isogenic wildtype (carrying native *tufA* and *tufB)* where relative fitness was set to 1.0. The relative fitness of each of the engineered constructs varied from 0.96 down to 0.77 (Figure 2). The relative fitness of *E. coli* in which *tufA* was deleted from the chromosome was 0.7. The similarity in relative fitness between *E. coli* lacking *tufA* and some of the strains carrying foreign *tuf* genes raised the question of whether all of the foreign genes would be capable of supporting viability in the absence of a functioning *tufB* gene.

**Figure 2.**
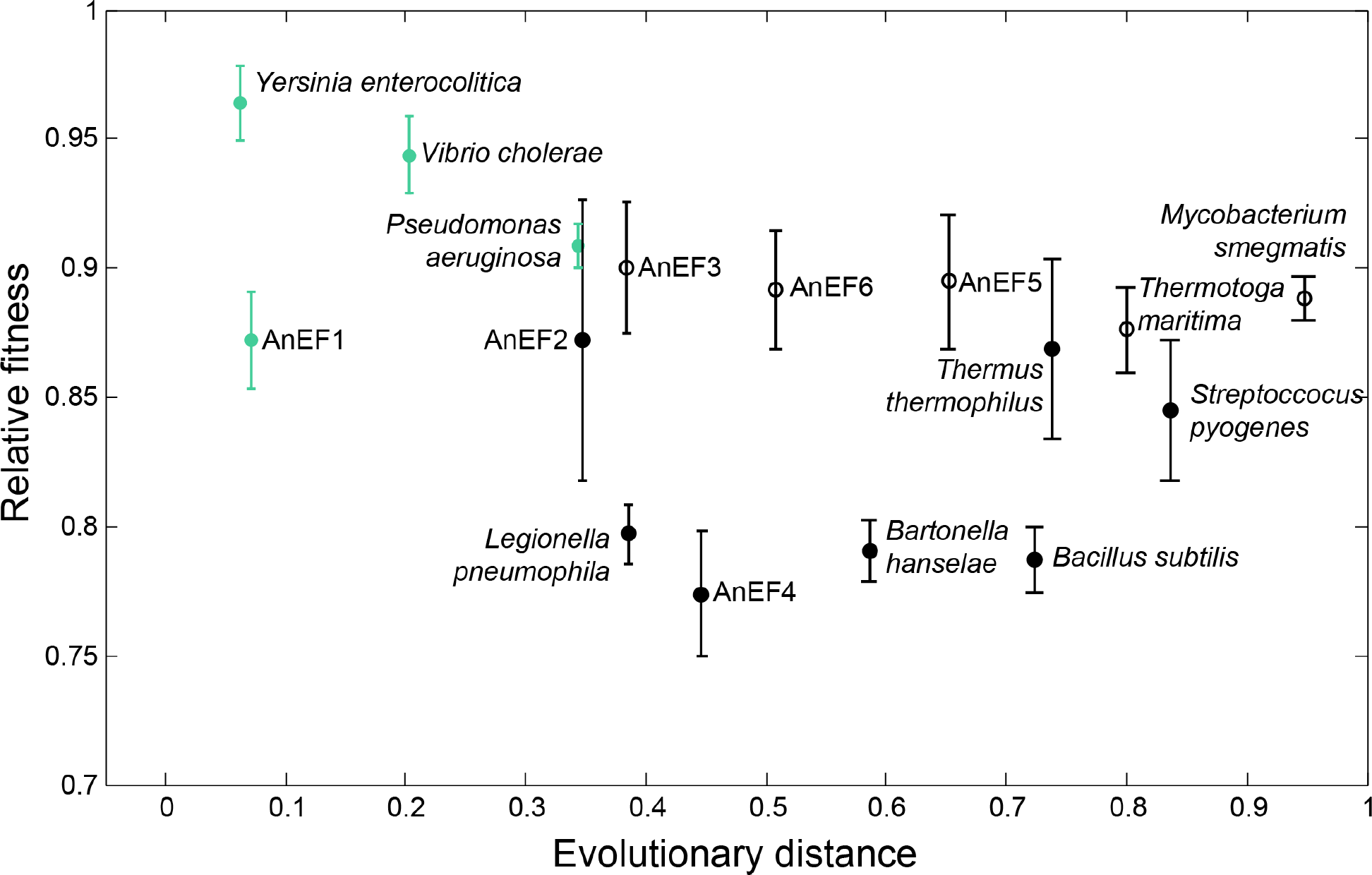
Correlation between relative fitness and evolutionary distance for bacterial strains carrying a foreign *tuf* gene. Relative fitness is shown as a function of evolutionary divergence (see Materials and Methods) of EF-Tu homologs from *E. coli* (1 indicates most different from *E. coli*). Each *E. coli* strain carried a foreign *tuf* gene at the *tufA* location and an intact native *tufB* gene. Fitness was measured as exponential growth rate, relative to *E. coli* wild-type carrying *tufA* and *tufB*. Species names and ancestor notations (AnEF, ancestral EF-Tu) refer to the source of the foreign *tuf* gene sequence in each strain. Strains shown in green carry a foreign *tuf* gene that can support viability even when *E. coli tufB* gene has been deleted. Strains shown in black carry foreign *tuf* genes that do not support viability in the absence of the *E. coli tufB* gene. Empty symbols represent strains where *tufB* region was amplified.

Because the *tufB* gene is located between the ribosomal RNA operons *rrnB* and *rrnE*, in a region that is subject to frequent amplification (Anderson and Roth 1981), we asked whether *tufB* was amplified in any of the strains carrying a foreign *tuf* allele. Our expectation was that a less effective, or completely inactive, foreign *tuf* gene, might select for genomes in which *tufB* was amplified as a fitness-compensatory mechanism. Using real-time quantitative PCR (RT-qPCR) we found that the *tufB* region was duplicated or triplicated in 5 of the 16 engineered strains (Figure 2). The 5 strains in which the *tufB* region was amplified are those that carry the most distant *tuf* homologs, compatible with selection for improved fitness.

### A subset of foreign *tuf* genes support viability

We next asked whether any of the foreign *tuf* genes could support viability in the absence of *E. coli tufB*, by attempting to remove the endogenous *tufB* gene from the chromosome of each of the 16 strains carrying a foreign *tuf* gene, as outlined in Figure S1B. In the absence of *tufB*, the only foreign *tuf* sequences that supported viability were those from *Y. enterocolitica, V. cholerae, P. aeruginosa*, and AnEF1, the youngest of the ancestral *tuf* genes, at ca. 0.7 bya, (Figure 2), whereas the phylogenetically more distant *tuf* genes did not. The relative fitness of each of the 4 viable strains was assessed by comparing their growth rates to that of the isogenic *E. coli* Δ*tufB* strain (carrying native *tufA*). The relative fitness values ranged from 0.95 (Y. *enterocolitica)*, down to 0.40 in the case of the *tuf* sequence from *P. aeruginosa* (Figure 3).

**Figure 3.**
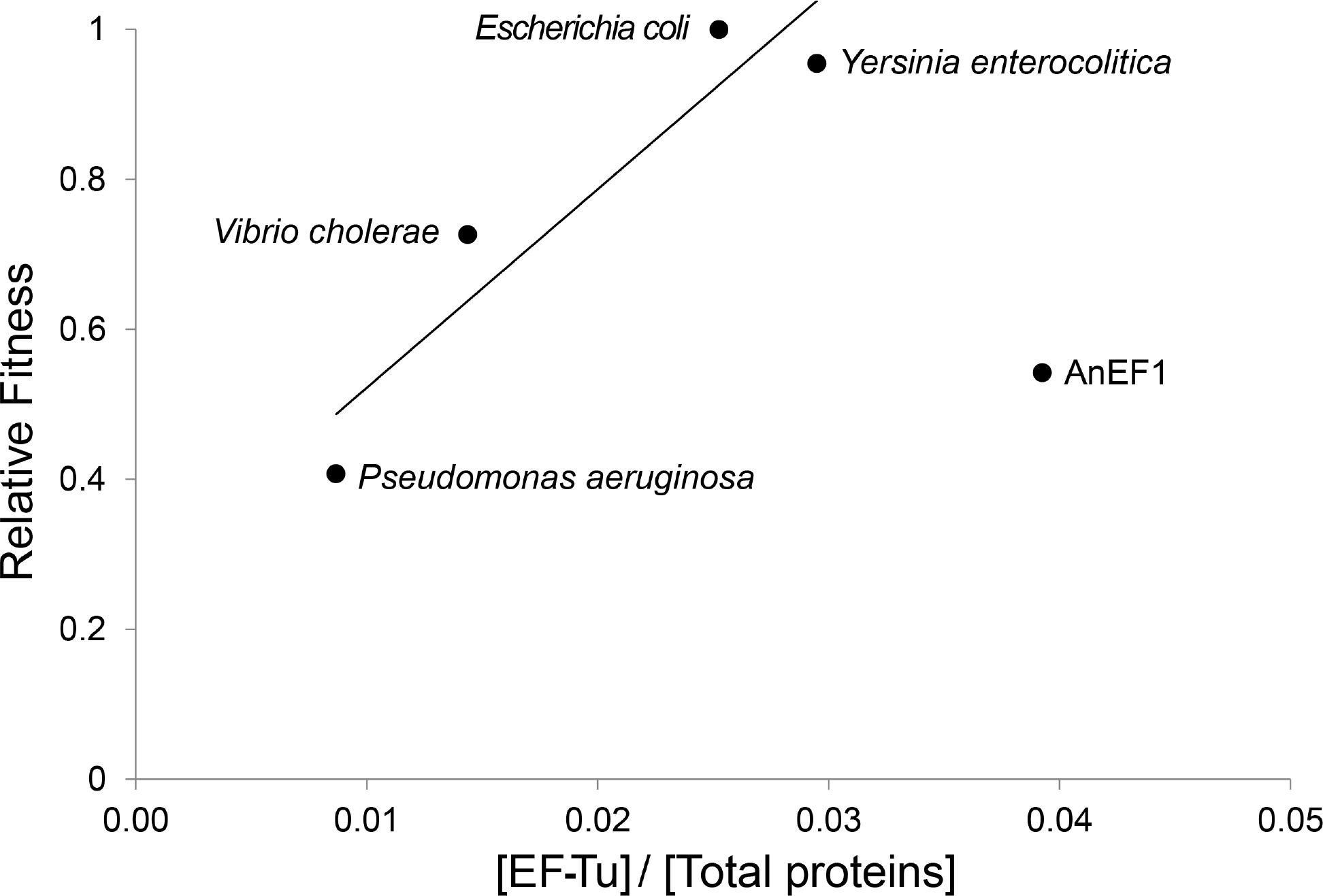
Relative fitness of strains carrying a single tuf gene, at the *tufA* locus, as a function of EF-Tu protein produced. EF-Tu concentration was normalized to the total protein concentration. Species names indicate the species origin of the only *tuf* gene present. (R=0.938).

### Fitness of engineered strains correlates with EF-Tu protein levels

A key question is why the EF-Tu gene replacements reduce fitness? Two possibilities (which are not mutually exclusive) are that the foreign genes are sub-optimally expressed (i.e. a concentration problem) or that they are sub-optimal in their function and interaction with the protein synthesis machinery and cellular network (i.e. an activity or toxicity problem). To evaluate the possibility that the foreign genes were sub-optimally expressed we measured EF-Tu abundance by LC-MS/MS in each of the viable strains (Figure 3). This analysis demonstrated that the level of EF-Tu relative to total protein varied significantly between engineered strains carrying different foreign *tuf* genes (Figure 3). With the exception of the ancestral gene, AnEFTU1, there is a good correlation between relative fitness and the concentration of EF-Tu. These data suggest that in most cases at least some of the reduction in relative fitness associated with foreign *tuf* genes is because bacteria do not produce an adequate level of EF-Tu to support fast growth. In the case of AnEFTU1 the reduced fitness may be more closely associated with reduced specific activity and/or toxicity.

### Conservation of EF-Tu residues correlates with viability in *E. coli*

The established function of EF-Tu is to deliver aminoacylated tRNAs into the A-site on mRNA-programmed ribosomes in order to drive rapid and accurate protein synthesis. This function requires that EF-Tu must have a sequence and structure that can efficiently interact with GTP, EF-Ts, each of the elongator tRNAs, and with the ribosome. EF-Tu has a highly conserved structure which must be capable of undergoing major structural rearrangements during its functional cycle (Nilsson and Nissen 2005; Yu, et al. 2009). All 17 EF-Tu sequences were aligned with the EF-Tu sequence from *E. coli* (Figure 4) to aid visualization of amino acids differences between EF-Tu’s associated with viability versus non-viability. The four viable foreign EF-Tu’s (green in Figure 4) show a bias towards a higher conservation of amino acid identity (84% to 94%), relative to *E. coli* EF-Tu, than the twelve non-viable EF-Tu’s (82% to 69%). It is interesting to note that there is only a small difference in total percent identity between several of the non-viable EF-Tu’s, (AnEF2, AnEF3, AnEF4, AnEF6, and *B. hanselae*, 80-82% similarity), and the most distantly related viable EF-Tu *(P. aeruginosa*, 84% similarity), suggesting that the loss of viability in these cases might be associated with changes in a few important residues of EF-Tu.

**Figure 4.**
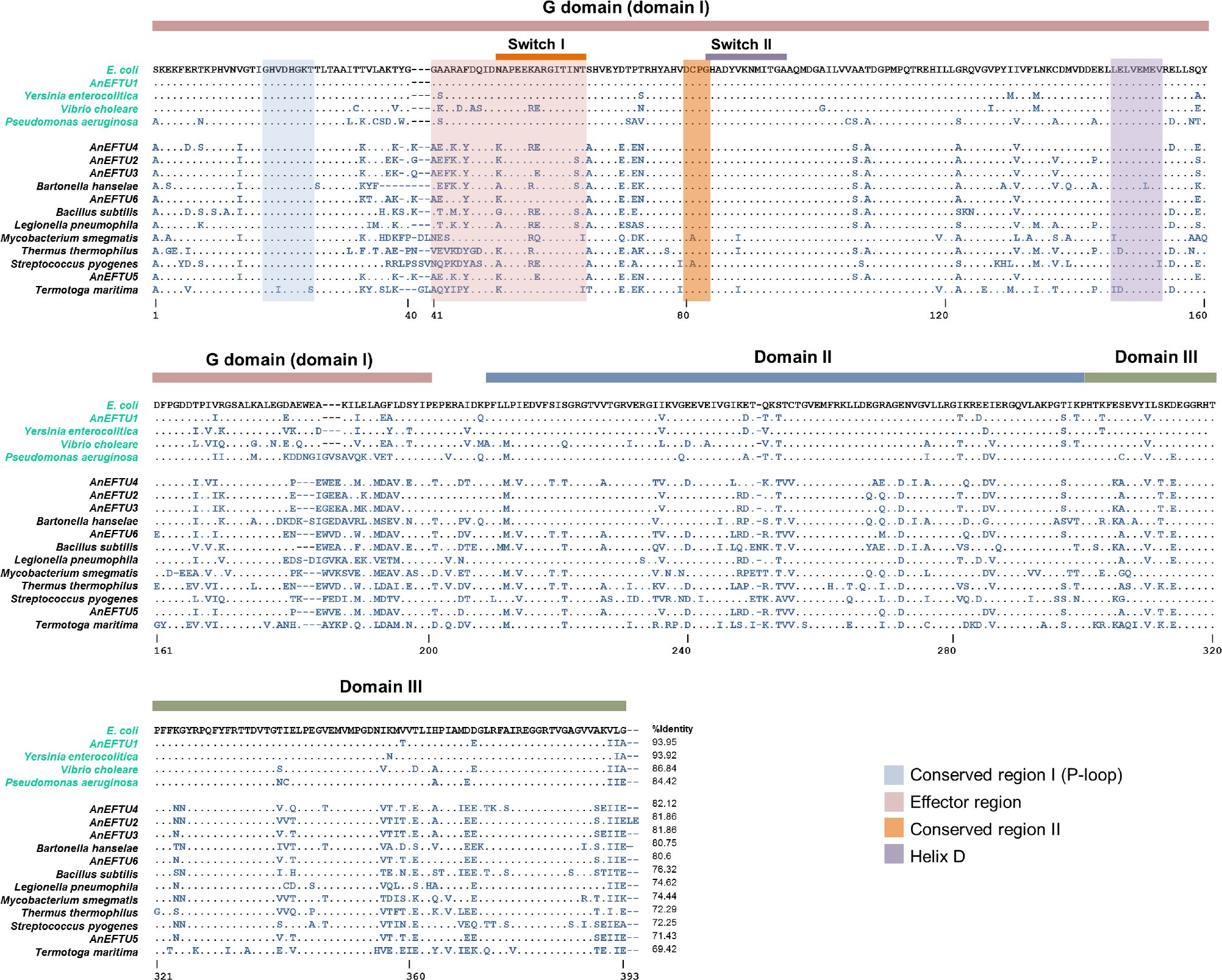
Alignment of EF-Tu protein sequences encoded by foreign *tuf* variants. EF-Tu sequences were aligned using Clustal W. Sequences labeled in green indicate that the foreign *tuf* gene supports viability as the only *tuf* gene in the genome. Sequences labeled in black indicate that the strain requires *E. coli tufB* for viability.

### Connectivity analysis of EF-Tu reveals an extensive interaction network

Several studies suggest that protein connectivity in a given network might modulate its activity in the cell (Fraser, et al. 2002; Hahn, et al. 2004; Lemos, et al. 2005). To examine this, we retrieved an *E. coli* interactome from the HitPredict database (Patil, et al. 2011) and measured the connectivity of EF-Tu in this network. The connectivity of EF-Tu was quantified by measuring its so-called ‘degree centrality’. Degree centrality is defined as the number of interactions that a given node has in its network. The average degree centrality of all proteins in the *E. coli* interactome is 12, whereas the degree centrality of EF-Tu is much higher at 172. Relative to all proteins in the interactome, EF-Tu ranks among the top ten most connected proteins in *E. coli* (Supplementary Table 1). Interestingly, the set of its first interaction partners is enriched in essential proteins (p-value = 3.66 × 10^-21^), as calculated via the HitPredict Database (Figure S4). Accordingly, this analysis suggests the possibility that, in addition to affecting the specific activity of EF-Tu in protein synthesis, additional deleterious effects on fitness might be due to the foreign *tuf* homologs disturbing the extensive interaction network of EF-Tu.

## Discussion

We generated *E. coli* strains in which the *E. coli tufA* gene was replaced by ancestral and modern homologs of *tuf*, from a broad spectrum of species and ancestral nodes. The origins of the ancestral EF-Tu sequences ranged in age from the Precambrian era, approximately 0.7 bya, back to the last universal common bacterial EF-Tu ancestor, approximately 3.6 bya (Figure 1). We showed that homologs of EF-Tu encoded by *tuf* genes from within the γ-proteobacteria, including one of the reconstructed ancestral node sequences, AnEF1, are functionally active in *E. coli* and support viability when present as the only *tuf* gene in the chromosome (shown in green in Figure 2). In contrast, more distantly related homologs and ancestral sequences were unable to support viability as the sole *tuf* gene. Among the four viable homologs, there was a good correlation (R = 0.938) between phylogenetic distance from *E. coli* EF-Tu and the magnitude of reduced growth fitness for the three homologs from extant species (Figure 2). The exception was the reconstructed ancestral node sequence, AnEF1, where the decrease in relative fitness was much greater than predicted by phylogenetic distance (Figure 2).

The correlation between phylogenetic distance from *E. coli* and relative growth fitness for the viable homologs raises the question of whether the underlying cause of the reduced fitness is a reduction in the specific activity of the foreign EF-Tu’s and/or a reduction in the amount of EF-Tu produced. By measuring EF-Tu protein concentration as a function of total protein concentration for each viable strain, including *E. coli* carrying only *tufA*, we observed a strong correlation between relative growth fitness and EF-Tu concentration for the each of the four EF-Tu’s from extant species (Figure 3). The slope of the correlation suggests that most of the reduced fitness associated with the EF-Tu homologs from the extant species could be attributed to a reduced level of EF-Tu rather than a reduced activity. Consistent with this, experiments have shown that reductions in growth rate are linearly correlated with reductions in cellular EF-Tu concentrations (Tubulekas and Hughes 1993b). Once again the exception to this correlation is found for the reconstructed ancestral AnEF1, where a very high concentration of EF-Tu was associated with a low relative fitness (Figure 3) consistent with AnEF1 having a low specific activity. This conclusion is supported by the results of *in vitro* translation, where in a system containing only *T. thermophilus* components (tRNA’s, EF-Ts, ribosomes) AnEF1 supported protein synthesis but with only 30% of the activity of native *E. coli* EF-Tu, demonstrating that AnEF1 can participate in peptide synthesis, albeit in a diminished fashion relative to their modern counterpart EF-Tu (Zhou, et al. 2012).

Why, given that the *tuf* gene regulatory regions are identical in all strains, do the foreign *tuf* coding sequences cause a reduction in the level of EF-Tu produced? Possible explanations include that the nucleotide sequence introduced with the foreign genes affects *tuf* mRNA halflife as previously shown for a mutant of *tuf* (Hammarlöf and Hughes 2008), or that the altered nucleotide sequence reduces translation efficiency, possibly by effects mediated through altered codon usage, or by affecting transcription-translation coupling, as recently shown for *tufB* (Brandis, et al. 2016; Brandis and Hughes 2016).

An important question is why some of the not-too-distant foreign homologs are unable to support viability. At the level of total amino acid similarity there is very little separating viable from non-viable EF-Tu sequences, with the boundary falling at approximately 84% similarity to *E. coli* EF-Tu (Figure S2). Given the very high level of conservation of EF-Tu it is possible that the non-viability of some foreign EF-Tu’s might be related to the alteration of just one or a few critically important residues. Indeed, many single amino acid substitutions in EF-Tu have been shown to generate protein variants that do not support viability (Abdulkarim, et al. 1991), including one single amino acid substitution in EF-Tu that permits ternary complex formation but abolishes translation activity by preventing ternary complex interaction with the ribosome (Tubulekas and Hughes 1993b). To facilitate an assessment of the amino acid differences between viable and non-viable EF-Tu’s their amino acid sequences were aligned (Figure 4). Of the 393 residues in *E. coli* EF-Tu, there were 116 residues that were identical in all of the viable EF-Tu homologs, but differed in at least one of the 12 non-viable homologs. Variation at one or more of these residues might explain the difference between EF-Tu viability and non-viability in *E. coli*. The 116 residues were distributed between each of the three structural domains of EF-Tu, with 48 in the G-domain, 38 in domain 2 and 30 in the C-terminal domain 3 (Figure S3-A). Several of the 116 variant residues lie within functionally important regions of EF-Tu including those involved in coordinating GTP hydrolysis, interaction with EF-Ts, and interaction with the ribosome (Kothe, et al. 2004; Kavaliauskas, et al. 2012; Thirup, et al. 2015). For example, in each of the non-viable EF-Tu’s there were from 3 – 8 variant residues in regions of the G-domain that are known to be functionally important for coordinating the hydrolysis of GTP on EF-Tu during protein synthesis. These variant residues are located in the Conserved regions I and II, the Effector region, and the Switch I and Switch II loops of the G-domain of EF-Tu (Figure S3-B). The altered residues potentially affecting GTP hydrolysis are at V20, G41, A43, R44, F46, N51, N63, T64, C81, and V88 (Figure 4). In addition, there are alterations in the P-loop, the Switch II region, and parts of domain 3 that are involved in interactions with EF-Ts and in helix D of EF-Tu (residues 144 – 156), that is involved in interactions with EF-Ts and protein L7/L12 on the ribosome (Kothe, et al. 2004; Thirup, et al. 2015). We can only speculate on the exact reason for the non-viability of these EF-Tu homologs. It seems unlikely that it is directly related to defects in binding or hydrolyzing GTP, given that this process involves highly conserved residues and structures and that the EF-Tu’s from extant organisms must be capable of supporting viability, including GTP binding and hydrolysis, in their natural system. Similarly, each of the ancestral homologs can support *in vitro* translation, albeit at a low efficiency, arguing that they also can bind and hydrolyze GTP. A similar line of reasoning could also rule out interactions with aa-tRNA’s as the cause of non-viability. Perhaps the most plausible reason for non-viability is defective interactions with EF-Ts and/or the ribosome. It seems reasonable to suggest that EF-Tu has co-evolved with EF-Ts and the ribosome, to modulate the efficiency of these interactions in each species. We suggest accordingly that the cause of non-viability for distantly related EF-Tu’s is not that they cannot function as enzymes capable of forming a ternary complex and hydrolyzing GTP, but rather that they are defective in one or more of the other important interactions made by EF-Tu, namely, with EF-Ts, with the mRNA-programmed ribosome, and possibly even interactions outside of protein synthesis involving one or more members of EF-Tu’s extensive protein interaction network (Figure S4). Co-evolution of EF-Tu with its interaction partners would create a barrier to transfer for EF-Tu’s beyond a certain threshold.

We previously hypothesized that evolutionary novelties are more likely to be shared between a descendant and its ancient homolog than between two currently existing protein homologs (Kacar and Gaucher 2012). Accordingly, replacing an existing gene with its ancient homolog may have a smaller negative fitness impact on the organism relative to exchanging the native gene with a currently existing homolog. However, functional divergence occurring through time could result in ancestral sequences being so maladapted to the new host cell that a functional organism is all but precluded (Copley 2003). This limitation does not apply only to ancestral genes and it has been suggested that as the number of nodes connecting a protein within its protein-protein interaction network increases, the probability that a protein could be successfully replaced with a homolog will decrease even if there is a functional equivalence between the endogenous gene and the homolog (Jain, et al. 1999). While a careful assessment of candidate ancestral protein properties prior to integration is helpful, in most cases, studying gene-triggered genomic perturbations experimentally through the integration of ancestral genes offers a valuable and complementary alternative to existing methodologies that use extant homologous proteins (Johnsen and Levin 2010; Acevedo-Rocha, et al. 2013; Pal, et al. 2014; Hobbs, et al. 2015).

How can we identify the specific historical constraints on replacement? We observe that only the ancient EF-Tu representing an ancestor within the γ-proteobacteria, AnEF1 (0.7 bya), and the modern EF-Tu homologs from extant γ-proteobacteria, are viable. In contrast, the last common ancestor of the α-, β-, and γ-proteobacteria, AnEF6 (1.3 bya) is non-viable. Accordingly, we speculate that mutational substitutions in EF-Tu occurring between 1.6 bya and 0.7 bya influenced the replaceability of *tuf* genes. These mutations may constrain *tuf* replaceability by disturbing the EF-Tu’s functional interaction with other cellular components, ultimately impacting its participation in protein synthesis. Thus, extensive mutational remodeling of interaction partners may be necessary in order to engineer even older ancient *tuf* genes inside the bacteria.

## Conclusions

We show that foreign *tuf* genes encoding EF-Tu proteins exhibit suboptimal functionality and reduced fitness when introduced into another host. The sub-optimality of the foreign *tuf* genes most likely results from disturbances in interactions directly important for protein synthesis, but suboptimal EF-Tu protein levels and disturbance of other potentially important interactions in the network of EF-Tu might also play a role. The observation that the only *tuf* homologs that supported viability belong to γ-proteobacterial taxon, or an associated ancestral node within the γ-proteobacteria, suggests that there is a relatively stringent “transferability cutoff” i.e., a point in the phylogeny beyond which functional divergence is too great for replacement. For EF-Tu protein this transferability zone is within the ancestral and modern γ-proteobacterial taxon, unlike some ribosomal proteins where constraints on replaceability are less stringent (Condon, et al. 1995; Lind, Tobin, et al. 2010).

Future efforts may involve identifying protein sites that interfere with organismal level function, and epistatically inhibit an ancient protein’s function in a descendant organism. Our experiments suggest that a protein like EF-Tu, that is highly conserved, and involved in multiple highly conserved interactions, is so highly optimized and fine-tuned in the host organism that it is essentially irreplaceable with distantly related foreign genes. The degree to which epistatic interactions constrain EF-Tu replaceability and functionality in the cell needs to be studied more to deepen our understanding of the design principles of complex biological systems and to allow us to introduce alterations in modern organisms by genetic engineering and gene replacements.

## Acknowledgments

Siv Andersson, Brian Hammer, Eric Gaucher and Lionel Guy provided the DNA sequences for some of the EF-Tu homologs used in this study. We thank BMSIS YSP undergraduate fellow Gokce Senger for her assistance with the curation of the EF-Tu sequences and Alison Cloutier for her assistance with the branchlength distance calculations. This work was supported by a NASA Astrobiology Postdoctoral Fellowship (BK), a research grant by the John Templeton Foundation (BK), grants from the Swedish Research Council (DH, DIA), and from the Knut and Alice Wallenberg Foundation, RiboCORE project (DIA & DH). The opinions expressed in this publication are those of the authors and do not necessarily reflect the views of any particular organization.

## Media and growth conditions

In general, liquid and solid media used were Luria Bertani (LB) media (10 g NaCl -5g in the case of low-salt LB-, 5 g yeast, 10 g triptone) and LA plates (LB with 1.5% of agar). For sacB counter-selections the LB and LA media used had no NaCl and LA media was also supplemented with sucrose 5 %. All incubations were made at 37°C (unless stated otherwise), and liquid cultures were shaken at 200rpm for aeration. The antibiotics used (Sigma-Aldrich, Sweden) had the following final concentrations: kanamycin 50 μg/mL and tetracyclin 15 μg/mL.

## Bacterial strains

All strains used in this study are derived from sequenced wild-type *E. coli* K12 strain MG1655 (Hayashi et al, 2006), unless stated otherwise. A list of the strains used in these experiments is shown in Table 1.

## Strain construction

The genetic marker [TP22-amilCP_opt-kan-sacB-T0], was inserted into the chromosome to replace either *tufA* or *tufB* by double-stranded DNA lambda-red recombineering (Datsenko & Wanner, 2000; Yu et al., 2000). The lambda-red genes were induced in each strain from the temperature-sensitive pSIM5-tet plasmid by incubation of an over-day culture (OD_600_: 0.3) at 43°C for 15 minutes. After cooling for 10 minutes in ice, the cells were made electrocompetent by washing in ice-cold water three times. Electroporation of [TP22-amilCP_opt-kan-sacB-T0] PCR product was done with a Gene Pulser (Bio-Rad, USA) by mixing 50 μl of electrocompetent cells and 100 ng of the cassette, with settings 1.8 kV, 25μF and 200 Ω. Cells were recovered in 1mL of low-salt LB at 30 °C overnight with agitation, and after recovery 100 μl of the culture were spread in LA plates containing kanamycin for selection of recombinants.

Genetic markers were moved between strains by phage-mediated (P1 *virA*) transduction. Lysates of the strains carrying the marker were made by mixing 1 mL of overnight culture containing 5mM CaCl_2_ with 100 μΙ of the P1 virA lysate previously made on E. coli MG1655. The bacteria:phage mix was incubated for 10 minutes and then 4mL of soft-agar (LB media + 0.8% agar + 5mM CaCl_2_) were added; this mixture was spread over an LA plate and incubated overnight. To release the bacteriophages the soft-agar was mixed with 4 mL and vortexed. The resultant slurry was centrifuged for 15 minutes at 5000 rpm and the supernatant was filtered through a 0.2 μm filter. Markers were transduced into the desired recipient strain by mixing 100 μl of the lysate with 500 μl of an overnight culture containing 5 mM CaCl_2_. After 10 minutes of incubation, 100 μl of the mixture was spread onto a selective plate.

Insertions of alien *tuf* genes were made by double-stranded DNA lambda-red recombineering to replace the previously inserted counter-selectable [TP22-amilCP_opt-kan-sacB-T0] marker. To make clean deletions, markers were deleted by single stranded lambda-red recombineering with counter selection for sucrose resistance (Ellis et al., 2001), following the same steps as commented above for the double-stranded DNA lambda-red recombineering.

## PCR and oligonucleotides

Polymerase Chain Reaction (PCR) was performed on a S1000™ Thermal Cycler (Bio-Rad, USA). Oligonucleotides were designed with the software CLC Main Workbench 7 (CLC bio, Denmark) using the genome of E. coli MG1655 as reference. For generation of the [TP22-amilCP_opt-kan-sacB-T0] cassette, PCR was performed using Phusion^®^ High-Fidelity PCR Master Mix with HF buffer (New England Biolabs, USA) and with the following cycling conditions: 98 °C for 30’’; 30 cycles of 98 °C for 10’’, 55°C for 30’’, 72 °C for 4’; 72 °C for 7’. For routine diagnostic PCR, Fermentas PCR Master Mix (Thermo Scientific, USA) was used with the following cycling conditions: 95 °C for 5’; 30 cycles of 95 °C for 30’’, annealing temperature (TA for 30’’, 72 °C for elongation time (E_T_)’; 72 °C for 5’. T_A_ varied depending on the pair of primers used and E_T_ was based on the length of the expected product (30’’ per kb). Oligonucleotides for construction of strains, PCR and sequencing are shown in Table S1.

## Preparation of genomic DNA and quantitative real-time PCR

Genomic DNA prepared using the MasterPure™ DNA Purification Kit (Epicentre Biotechnologies, USA) was used to run quantitative real-time PCR. 1 μΕ gDNA (diluted 1:10, 1:100, 1:1000 and 1:10000), 10 *μΕ* PerfeCTa SYBR Green FastMix (Quanta BioSciences), 0.6 μΕ of 10 μΜ forward and reverse primers, and ddH2O were added to a final reaction volume of 20 μυ The Eco Real-Time PCR Systems (Illumina) was used for running the PCR. Amplification of the cysG and indT genes was used as standard. The oligonucleotides used as qRT-PCR primers are listed in Table S1.

## Whole genome sequencing and analysis

Genomic DNA was prepared using the MasterPure™ DNA Purification Kit (Epicentre Biotechnologies, USA). To create libraries of paired-end fragments, Nextera XT Sample Preparation Kit (Illumina, USA) was used following instructions from manufacturer. Sequencing was performed on the Illumina MiSeq, generating 250 bp paired-end reads. Whole genome sequencing data was analyzed using the software CLC Genomic Workbench (CLC bio, Denmark)

## Local DNA sequencing

Local sequencing of PCR amplified products was performed at Macrogen Europe sequencing facilities (Amsterdam, The Netherlands), and data was analyzed using the software CLC Main Workbench 7 (CLC bio, Denmark).

## Growth rates and relative fitness measurements

Growth rate was measured by monitoring the rate of increase in optical density using a Bioscreen C machine (Oy Growth Curves Ab Ltd, Findland), growing the cultures in a Honeycomb microtiter plate. To set up an experiment, 0.5 μl of overnight culture from a single colony was diluted into 1mL of LB medium. For each independent colony, 3 wells of the microtiter plate were filled with 300 μl of each dilution, as technical replicates. Cultures were grown for 18h at 37 °C with continuous shaking and readings of OD_600_ were taken at 5 minutes intervals. The doubling time (dt) of each strain during exponential growth was estimated over an interval of 50 minutes beginning at OD_600_=0.025. Relative fitness was determined in each case by comparing the doubling time to the average of all the wild-type measurements.

## Statistic analysis

All statistical analyses were performed using GraphPad Prism® v6.0c (GraphPad Software, USA). The significance of differences between fitness costs was calculated using an unpaired two-tailed t-test.

## Proteomics

**Cell lysis, digestion and labeling procedures.** All cells were placed into Covaris® microTUBE-15 (Woburn, MA) microtubes with Covaris® TPP buffer. Samples were lysed in an Covaris S220 Focused-ultrasonicator instrument with 125W power over 180s with 10% max peak power. Lysed cells were digested via FASP digest according to the FASP Filter Aided Sample Prep protocol for trypsin digestion, followed by HPLC purification. We used Promega® Sequencing Grade Trypsin/LysC (V5073, Madison, WI) overnight at 38°C. Each sample was submitted for a single LC-MS/MS experiment that was performed on a LTQ Orbitrap Elite (Thermo Fischer) equipped with a Waters (Milford, MA) NanoAcquity HPLC pump or Orbitrap Lumos (Thermo Fischer, San Jose, CA) equipped with EASYLC1000 (Thermo Fischer, San Jose, CA). Peptides were separated onto a 100 μm inner diameter microcapillary trapping column packed first with approximately 5 cm of C18 Reprosil resin (5 μm, 100 Å, Dr. Maisch GmbH, Germany) followed by analytical column ∼20 cm of Reprosil resin (1.8 μm, 200 Å, Dr. Maisch GmbH, Germany). Separation was achieved by applying a gradient from 5–27% ACN in 0.1% formic acid over 90 min at 200 nl min-1. Electrospray ionization was enabled by applying a voltage of 1.8 kV using a home-made electrode junction at the end of the microcapillary column and sprayed from fused silica pico tips (New Objective, MA). The LTQ Orbitrap Elite/Lumos was operated in data-dependent mode for the mass spectrometry methods. The mass spectrometry survey scan was performed in the Orbitrap in the range of 395–1,800 m/z at a resolution of 6 × 10^4^, followed by the selection of the twenty most intense ions (TOP20) for CID-MS2 fragmentation in the Ion trap using a precursor isolation width window of 2 m/z, AGC setting of 10,000, and a maximum ion accumulation of 200 ms. Singly charged ion species were not subjected to CID fragmentation. Normalized collision energy was set to 35 V and an activation time of 10 ms. Ions in a 10 ppm m/z window around ions selected for MS2 were excluded from further selection for fragmentation for 60 s. The same TOP20 ions were subjected to HCD MS2 event in Orbitrap part of the instrument. The fragment ion isolation width was set to 0.7 m/z, AGC was set to 50,000, the maximum ion time was 200 ms, normalized collision energy was set to 27V and an activation time of 1 ms for each HCD MS2 scan.

## Mass spectrometry analysis

Raw data were submitted for analysis in Proteome Discoverer 2.1.0.81 (Thermo Scientific) software. Assignment of MS/MS spectra was performed using the Sequest HT algorithm by searching the data against a protein sequence database including all entries from the users database, and our *E. coli* K12 database as well as other known contaminants such as human keratins and common lab contaminants. Sequest HT searches were performed using a 20 ppm precursor ion tolerance and requiring each peptides N-/C termini to adhere with Trypsin protease specificity, while allowing up to two missed cleavages. 6-plex TMT tags on peptide N termini and lysine residues (+229.162932 Da) was set as static modifications while methionine oxidation (+15.99492 Da) was set as variable modification. A MS2 spectra assignment false discovery rate (FDR) of 1% on protein level was achieved by applying the target-decoy database search. Filtering was performed using a Percolator 64 bit. For quantification, a 0.02 m/z window centered on the theoretical m/z value of each the six reporter ions and the intensity of the signal closest to the theoretical m/z value was recorded. Reporter ion intensities were exported in result file of Proteome Discoverer 2.1 search engine as an excel tables.

## Evolutionary Divergence

Two independent approaches were utilized in order to estimate the evolutionary distance between sequences, both leading the same evolutionary distance estimate output. i) MEGA software: The analyses involved 18 amino acid sequences. Analyses were conducted using the Poisson correction model (Zuckerkandl 1965). All positions containing gaps and missing data were eliminated. There were a total of 385 positions in the final dataset. Evolutionary analyses were conducted in MEGA7 (Kumar, et al. 2016). ii) The branchlenght distances were also calculated via a custom script that used ETE Software (Huerta-Cepas, et al. 2010). The custom python ETE v. 3 python library script is provided under supporting information.

## Computational methods

Computational analyses include i) the pairwise and multiple sequence alignments, ii) protein structure analysis, and iii) protein interaction network analysis. Clustal Omega was used to perform multiple sequence alignment with default parameters. Pairwise sequence alignments of the wild type EF-Tu against each ancient and modern EF-Tu proteins were performed by EMBOSS Needle of Clustal Omega with default parameters which uses the Needleman-Wunsch alignment algorithm.

The protein interactome of *E. coli* was retrieved from the HitPredict database and the network analysis was performed with the network package of Python. The sub-network of EF-Tu interaction partners was visualized with Cytoscape.

